# Predicting Endometrial Cancer Subtypes and Molecular Features from Histopathology Images Using Multi-resolution Deep Learning Models

**DOI:** 10.1101/2020.02.25.965038

**Authors:** Runyu Hong, Wenke Liu, Deborah DeLair, Narges Razavian, David Fenyö

**Affiliations:** Institute for Systems Genetics, NYU Grossman School of Medicine, New York, NY 10016, USA; Department of Biochemistry and Molecular Pharmacology, NYU Grossman School of Medicine, New York, NY 10016, USA; Department of Pathology, NYU Langone Health, New York, NY 10016, USA; Department of Population Health, NYU Langone Health, New York, NY 10016, USA

**Keywords:** endometrial carcinoma, cancer imaging, deep learning, computational pathology, computational biology, cancer genomics

## Abstract

The determination of endometrial carcinoma histological subtypes, molecular subtypes, and mutation status is a critical diagnostic process that directly affects patients’ prognosis and treatment options. Compared to the histopathological approach, however, the availability of molecular subtyping is limited as it can only be accurately obtained by genomic sequencing, which may be cost prohibitive. Here, we implemented a customized multi-resolution deep convolutional neural network, Panoptes, that predicts not only the histological subtypes, but also molecular subtypes and 18 common gene mutations based on digitized H&E stained pathological images. The model achieved high accuracy and generalized well on independent datasets. Our results suggest that Panoptes has potential clinical application of helping pathologists determine molecular subtypes and mutations of endometrial carcinoma without sequencing.

**Significance:** Recently, molecular subtyping and mutation status are increasingly utilized in clinical practice as they offer better-informed prognosis and the possibility of individualized therapies for endometrial carcinoma patients. Taking advantage of the multi-resolution nature of the whole slide digital histopathology images, our Panoptes models integrate features of different magnification and make accurate predictions of histological subtypes, molecular subtypes, and key mutations in much faster workflows compared to conventional sequencing-based analyses. Feature extraction and visualization revealed that the model relied on human-interpretable patterns. Overall, our multi-resolution deep learning model is capable of assisting pathologists determine molecular subtypes of endometrial carcinoma, which can potentially accelerate diagnosis process.

## Introduction

Endometrial cancer is the most common type of gynecologic cancer among women around the world with a rising occurrence and mortality (Amant et al., 2005; Burke et al., 2014a, 2014b; Morice et al., 2016). In the United States, it is one of the top 5 leading cancer types with 52,600 new cases reported in 2014. This number increased to 60,050 in the year of 2016, and was estimated to further increase to 61,880 in 2019 (Burke et al., 2014a, 2014b; Siegel et al., 2016). Globally, endometrial cancer caused approximately 42,000 women’s death in 2005, and this annual mortality count estimate drastically increased to 76,000 in 2016 (Amant et al., 2005; Morice et al., 2016). The 5-year survival rate, depending on the study cohort, is ranging from 74% to 91% for patients without metastasis (Burke et al., 2014a).

Clinically, endometrial carcinomas are stratified based on their grade, stage, hormone receptor expression, and histological characteristics (Bokhman, 1983). Histological classification reflects tumor cell type and informs the choice of surgical procedure and adjuvant therapy. The majority of endometrial cancer cases exhibit either endometrioid or serous characteristics, which comprise approximately 70%-80% and about 10% of all cases, respectively (Murali et al., 2014). Statistically, patients with serous subtype tumors have a lower 5-year survival rate due to more frequent metastases and a higher risk of recurrence (Morice et al., 2016). Thus, it is critical to determine the subtypes in order to determine patients’ individualized treatment plans and to assess prognosis (Amant et al., 2005; Frumovitz et al., 2004). Histological subtype is determined by pathologists after thorough examination of hematoxylin and eosin (H&E) stained tissue sample slides of tumor samples. Endometrioid tumors typically exhibit a glandular growth pattern, while the serous subtype is characterized by the frequent presence of a complex papillary pattern (Darvishian et al., 2004; Murali et al., 2019; Murray et al., 2000). These features are not exclusive for either of the subtypes, however, making histological classification challenging, especially among high grade cases, even for experienced pathologists and necessitating ancillary subtyping criteria (Brinton et al., 2013; Getz et al., 2013; Morice et al., 2016; Zannoni et al., 2010).

The multi-omics study of The Cancer Genome Atlas (TCGA) introduces a set of novel criteria that classify endometrial carcinoma into four molecular subtypes, namely POLE ultra-mutated, high microsatellite instability (MSI-high) hypermutated, copy-number low (CNV-L), and copy-number high (CNV-H), based on their mutation characteristics, copy number alterations, and microsatellite instability. This molecular classification standard has been gaining popularity among pathologists and clinicians in recent years. Among these four subtypes, patients with the CNV-H subtype, which includes serous carcinomas and a subset of high grade endometrioid cancers, had the worst outcomes based on progression free survival (Getz et al., 2013). Exome sequencing also revealed a panel of genes differentially mutated across the four molecular subtypes, many of which have been shown to play significant roles in endometrial carcinoma tumorigenesis and proliferation and can potentially be novel targets of individualized therapies (Bell, 2014; Liang and Lu, 2018). For example, most patients in CNV-H subtype are *TP53* mutated but *PTEN* wild-type. Determining the molecular subtyping and single gene mutations can provide new insights that complement and refine the histological classification, but the availability of this information is limited by the time and cost of sequencing.

New powerful computational approaches for analyzing massive biomedical data have tackled numerous challenges, which accelerates the pace of human health improvement worldwide. In particular, computational pathology, a discipline that involves the application of image processing techniques to pathological data, has been especially benefitted from the advancement of deep learning in recent years (Komura and Ishikawa, 2018; Louis et al., 2016; Madabhushi and Lee, 2016; Nawaz and Yuan, 2016). Convolutional neural network (CNN) models are capable of segmenting cells in histopathology slides and classifying them into different types based on their morphology (Cooper et al., 2018; Madabhushi and Lee, 2016). An InceptionV3-based model achieves a high level of accuracy in determining melanoma possibility, exhibiting significant diagnostic potential (Louis et al., 2016). Moreover, successful deep learning models have also been built to predict molecular and genomic features in cancer, such as microsatellite instability (MSI), immune subtypes, and somatic mutation status, suggesting that machine learning techniques may be able to assist human experts to further exploit clinically relevant information in pathological images (Coudray et al., 2018; Gillette et al., 2020; Hong et al., 2020; Kather et al., 2019; Kim et al., 2019; Wang et al., 2021). In addition, studies have also demonstrated that deep neural network models have the potentials in capturing features across cancer and tissue types (Fu et al., 2020; Kather et al., 2020).

H&E slides are often scanned at different resolutions and are saved into a single image file. This allows pathologists to examine features of various sizes at the optimal resolution. Here, we designed a customized architecture, that we call Panoptes. Panoptes takes advantage of the multi-resolution structure of the H&E image files. We showed that models using this architecture could classify common endometrial carcinoma histological subtypes, molecular subtypes, and several critical mutations with decent performance based on H&E images and outcompete existing InceptionResnet models in most top-performing tasks. Using tSNE dimensionality reduction technique, we extracted and visualized the features learned by models to classify H&E images. These histopathological features were mostly human interpretable, suggesting possibilities of incorporating them into the pathological diagnostic standards. In particular, we confirmed that tumor grade was the major factor to distinguish CNV-H molecular subtype from the other 3 molecular subtypes in the histological endometrioid cases. In addition, the generalizability of models was validated by independent datasets. By testing trained models of key predictive tasks on samples from NYU hospitals, their potential clinical capability was demonstrated.

## Results

### Data preparation and multi-resolution deep-learning based histopathology image analysis

The goal of this study was to build multi-resolution deep convolutional neural network models that could automatically analyze endometrial cancer digital H&E slides and predict their histological and molecular features. We used diagnostic formalin-fixed paraffin-embedded (FFPE) and H&E stained tumor slides and labels from 2 public datasets, TCGA and Clinical Proteomic Tumor Analysis Consortium (CPTAC), to train, validate, and test our models. TCGA and CPTAC are two mutually independent cohorts. Later, an independent dataset from samples at NYU hospitals were used to test the generalizability and potential clinical capability of select promising trained models. CPTAC cohort has 107 slides from 98 patients and TCGA cohort has 389 slides from 358 patients. Overall, 496 slides from 456 patients, covered in previous publications (Dou et al., 2020; Getz et al., 2013) and annotated with subtype and gene mutation information, were included to form a mixed TCGA-CPTAC dataset (Figure 1A and Figure S1A). More than 90% of patients in our cohort had only 1 diagnostic slide (Figure S1B). As a lot of driver gene mutations in endometrial cancer are correlated with histological and molecular subtypes, we validated these correlations to ensure that our cohort was a representative of the patient’s population (Figure S1C). The general process of training, validation, testing, and visualization followed the workflow in Figure 1B. For each prediction task, cases in the mixed dataset were randomly split into training, validation, and test set such that slides from the same patient were in only one of these sets. This allowed the test set to be strictly independent to the training process and also made it possible to obtain per-patient level metrics, which could be more useful in the clinical setting. Each task was performed on a different random split of cases stratified with the outcome. Promising models were selected based on their statistical metrics and further tested by the independent NYU dataset. Due to the extremely large dimension of the digital H&E slides (Figure S1D), slides were tiled into 299-by-299-pixel pieces and were packaged into one TFrecords file for each set after color normalization. All models were trained from scratch.

**Figure 1.**
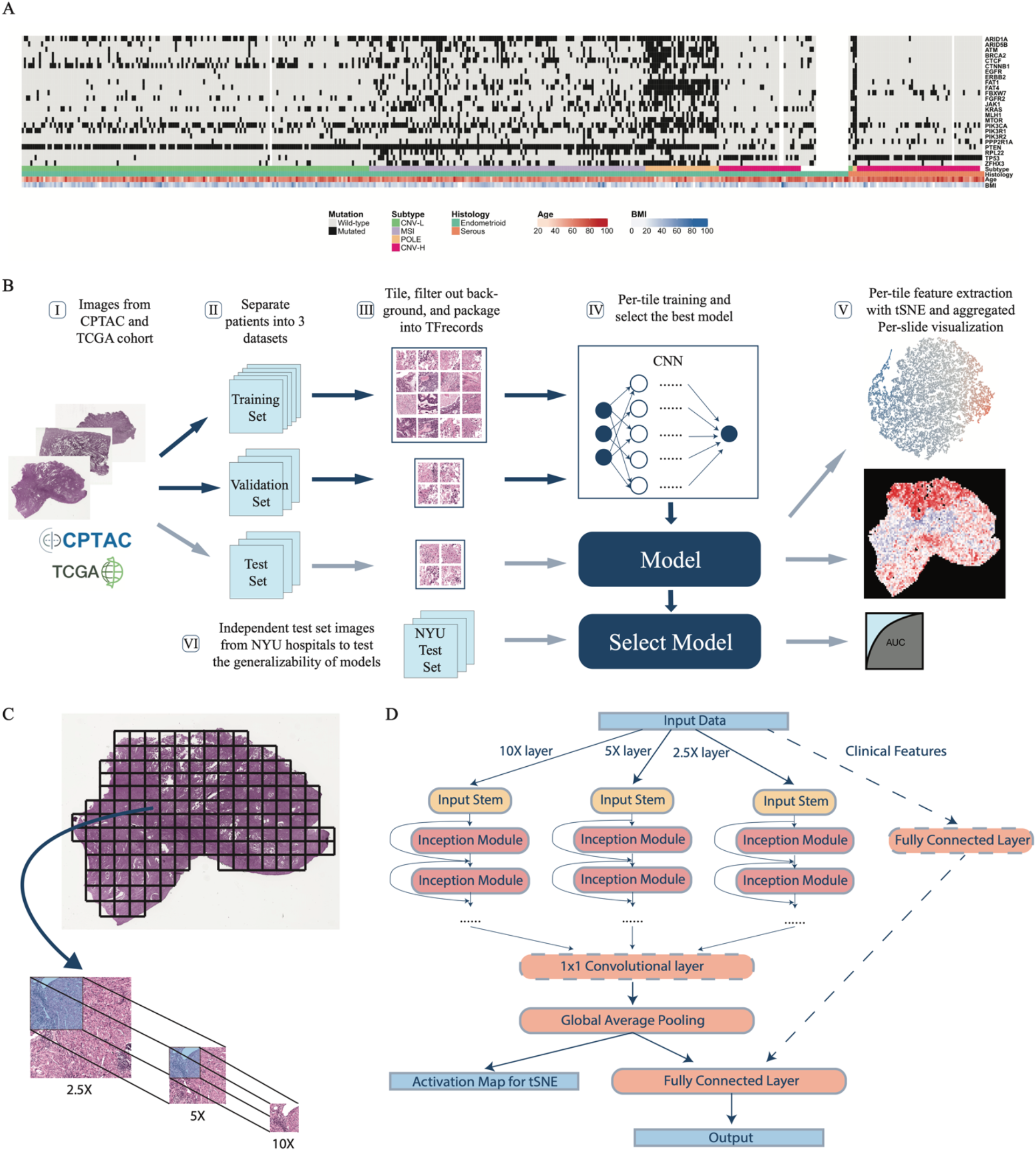
Workflow and Panoptes architecture. (A) Patients in the cohorts with feature annotations. (B) Overall workflow. I, H&E slide images of endometrial cancers were downloaded from databases; II, slides were separated at per-patient level into a training, a validation, and a test set; III, slides were cut into 299×299-pixel tiles excluding background and contaminants and qualified tiles were packaged into TFrecord files for each set; IV, training and validation sets were used to train the convolutional neural networks and the testing set was used to evaluate trained models; V, activation maps of test set tiles were extracted and dimensionally reduced by tSNE to visualize features while the per-tile predictions were aggregated back into intact slides. VI, an independent test set with samples from NYU hospitals was used to test the generalizability of selected best performing models. (C) Slides were cut into paired tile sets at 2.5X, 5X, and 10X equivalent resolution of the same region to prepare for Panoptes. (D) Panoptes architecture with optional 1×1 convolutional layer and clinical features branch.

We developed a multi-resolution InceptionResnet-based (Szegedy et al., 2017) convolutional neural network architecture, Panoptes, to capture features of various sizes on the H&E slides, which resembles the reviewing strategy of human pathologists. Unlike the conventional CNN architecture, the input of Panoptes is a set of 3 tiles of the same region on the H&E slide instead of a single tile. The equivalent scanning resolution of tiles in a set is 2.5X, 5X, and 10X so that the higher resolution tile covers one fourth of the region in the next lower resolution tile (Figure 1C). Hence, each grid region at 2.5X resolution can ideally generate 16 tile sets provided none of the tiles contained more than 40% of background pixels (Figure 1C). Each set of tiles were converted into a single matrix as one sample. Panoptes has three InceptionResnet-based branches, each of which processed the samples with a specific resolution of the same region simultaneously (Figure 1D). These branches worked separately until the third-to-last layer of the architecture, where inputs from the three branches were concatenated, followed by a global average pooling layer and the final fully connected layer. This design enabled the branches to learn features of different scales. More abstract information from each branch was integrated only at higher levels. We attempted to add an additional 1-by-1 feature pooling convolutional layer before the global average pooling and introduced a fourth branch processing clinical features, including patients’ age and body mass index (BMI). The effectiveness and comparisons of these modifications are discussed in later sections. Compared to conventional CNN, the multi-resolution design of Panoptes can at the same time considers both macro tissue-level features and minute cellular-level features of the same region, and can therefore capture more comprehensive characteristics of the slides. Moreover, taking the multi-resolution tile sets as input while having a single output and loss function preserves the original spatial information, which makes Panoptes distinct from the simply joining decisions from three separate models trained on tiles of three resolutions. To find the best performing model for different prediction tasks, we tried four different Panoptes architectures with and without the clinical feature branch, two types of InceptionResnet, and three types of Inception in this study. InceptionResnet and Inception models were trained on single resolution 10X tiles. Among all the statistical metrics calculated, we used area under the receiver operating characteristic curve (AUROC) of the test sets as the major metrics to evaluate the performance of the models, which is the typical standard in the machine learning field. Precision, recall, sensitivity, specificity, and F1 scores were also considered to evaluate imbalanced prediction tasks. Per-patient level prediction was obtained by taking the mean of the predicted probability (prediction score) of all tiles or tile sets belonging to the same patient. The AUROC was then calculated by taking each patient as one sample point. For Panoptes models, one set of tiles was counted as a single tile for the metric calculation purpose since the output from the model was only one prediction score for each set.

### Multi-resolution deep-learning architectures achieved better predictive performance on histopathology images

We trained models to predict histological subtypes, CNV-H subtype from the entire cohort and the endometrioid patients, CNV-L subtype, MSI-high subtype, POLE subtype, and the mutation status of 18 endometrial-carcinoma-related genes. We applied five baseline models (InceptionV1, InceptionV2, InceptionV3, InceptionResnetV1, and InceptionResnetV2) and four versions of multi-resolution models (Panoptes1-4) on all of the tasks (Figure S2A, S2B). The same data splits were used for all the models of the same predictive tasks in order to have fair comparisons among models with different architectures. The best performing architectures for each of the prediction tasks and their corresponding AUROC with 95% confidence intervals (CI) are shown in Table 1. Tasks with per-patient AUROC less than 0.6 were not listed. We performed 1-tail Wilcoxon tests on prediction scores between positively and negatively labeled tiles for the results in Table 1, which all showed significant differences (Figure 2A). Therefore, the prediction scores of true-label-positive tiles were significantly higher than those of true-label-negative tiles, demonstrating that these models were able to distinguish tiles in the test sets.

**Table 1.** AUROC of the best models for each task with 95% confidence intervals (CIs).

|  | **BEST**<br>**ARCHITECTURE** | **PER-PATIENT**<br>**AUROC** | **PER-TILE AUROC** |
| --- | --- | --- | --- |
| **HISTOLOGY** | Panoptes2 | **0.969 (0.905-1)** | **0.870 (0.866-0.874)** |
| **CNV-H FROM**<br>**ENDOMETRIOID** | Panoptes1 | **0.958 (0.886-1)** | **0.864 (0.859-0.870)** |
| **CNV-H** | Panoptes4 | **0.934 (0.851-1)** | 0.731 (0.728-0.734) |
| **POLE** | Multi-model system | **0.890 (0.821-0.960)** | 0.691 (0.683-0.700) |
| **CNV-L** | Panoptes1 | **0.889 (0.755-1)** | 0.710 (0.705-0.716) |
| **TP53** | Panoptes2 | **0.873 (0.768-0.977)** | 0.713 (0.709-0.717) |
| **FAT1** | Panoptes2 with<br>clinical | **0.835 (0.666-1)** | 0.639 (0.635-0.642) |
| **MSI-HIGH** | InceptionResnetV1 | **0.827 (0.705-0.948)** | 0.638 (0.635-0.641) |
| **ZFHX3** | InceptionResnetV1 | **0.824 (0.689-0.959)** | 0.637 (0.634-0.640) |
| **PTEN** | InceptionV2 | **0.781 (0.579-0.984)** | 0.623 (0.620-0.627) |
| **FGFR2** | Panoptes4 with<br>clinical | **0.755 (0.540-0.970)** | 0.550 (0.545-0.554) |
| **MTOR** | Panoptes1 | 0.724 (0.496-0.951) | 0.674 (0.670-0.678) |
| **CTCF** | Panoptes4 | 0.724 (0.518-0.931) | 0.571 (0.568-0.575) |
| **PIK3R1** | InceptionResnetV1 | 0.702 (0.524-0.880) | 0.596 (0.593-0.599) |
| **PIK3CA** | Panoptes4 | 0.689 (0.532-0.847) | 0.526 (0.523-0.530) |
| **ARID1A** | InceptionResnetV2 | 0.683 (0.513-0.853) | 0.542 (0.538-0.545) |
| **JAK1** | Panoptes2 with<br>clinical | 0.662 (0.410-0.940) | 0.612 (0.605-0.618) |
| **CTNNB1** | InceptionResnetV2 | 0.648 (0.439-0.858) | 0.619 (0.616-0.622) |
| **KRAS** | Panoptes2 with<br>clinical | 0.638 (0.404-0.871) | 0.515 (0.510-0.519) |
| **FBXW7** | InceptionV3 | 0.629 (0.366-0.892) | 0.606 (0.602-0.609) |
| **RPL22** | InceptionV3 | 0.632 (0.395-0.868) | 0.517 (0.512-0.522) |
| **BRCA2** | InceptionResnetV1 | 0.613 (0.318-0.908) | 0.624 (0.620-0.629) |

**Figure 2.**
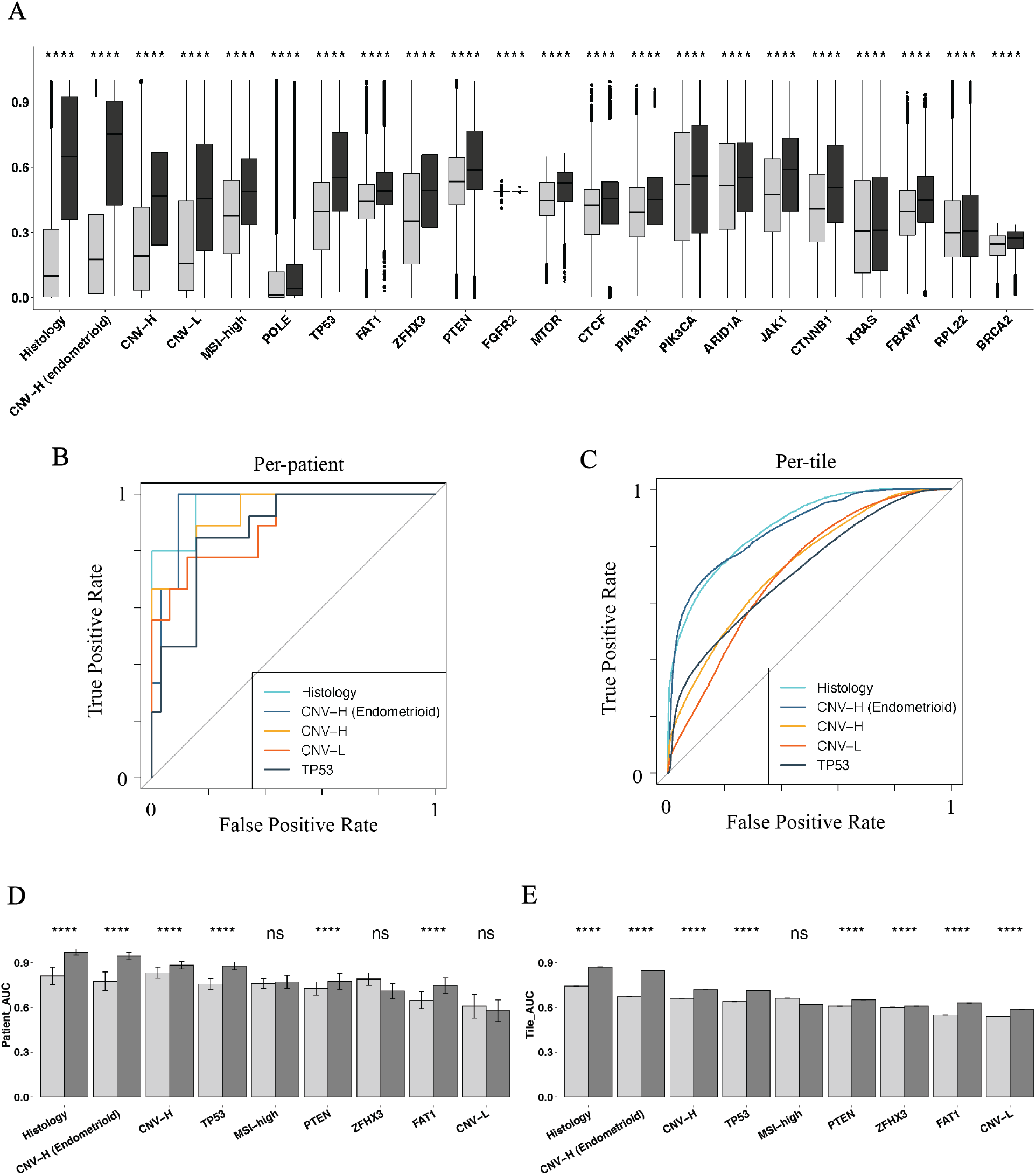
Prediction tasks were statistically successful with promising results and Panoptes outcompeted baselines in most of top-performing prediction tasks. (A) Predicted positive probability of tiles with 1-tail Wilcoxon test between true label positive and negative groups (black: true label positive tiles; grey: true label negative tiles) from models in **Table 1**. (B, C) ROC curves at per-patient (B) and per-tile (C) level associated with the top five prediction tasks in (A). (D, E) Bootstrapped per-patient (D) and per-tile (E) AUROC of InceptionResnetV2 (light) and Panoptes2 (dark) of top nine tasks in (A) with 1-tail t-test.

Based on the AUROC scores, we observed that Panoptes models were the best architectures in the six out of the top seven prediction tasks (Table1, and Figure 2B, 2C). It is also observed that Panoptes performed better than Inception and InceptionResnet models for most of the tasks (Figure S2A, S2B). To validate that Panoptes performed better than InceptionResnet, we conducted 1-tail t-test on AUROC performance of the top nine prediction tasks between the Panoptes models and their corresponding InceptionResnet models. Panoptes2, which was the best Panoptes architecture in most of the tasks, showed a significantly higher AUROC than the corresponding InceptionResnet2 in six prediction tasks at per-patient level and eight at per-tile level (Figure 2D, 2E). Similarly, Panoptes1 had a significantly higher AUROC than InceptionResnet1 in five prediction tasks at per-patient level and eight at per-tile level (Figure S3A, S3B).

To evaluate the effectiveness of adding an additional 1-by-1 convolutional layer between concatenation of branches and the global average pooling, we performed a 1-tail t-test between Panoptes1 and Panoptes3 as well as Panoptes2 and Panoptes4. However, only four tasks at per-patient level and six tasks at per-tile level showed significant p-values between Panoptes2 and Panoptes4 (Figure S3C, S3D). Similar results were observed between Panoptes1 and Panoptes3, where only one per-patient level task and four per-tile level tasks showed a significant difference. By applying the same test to Panoptes with and without clinical feature branch models, most of the tasks were not statistically significant with an example of Panoptes2 having four significant tasks at per-patient level and five at per-tile level (Figure S3E, S3F). In summary, our multi-resolution architectures Panoptes outperformed InceptionResnet in analyzing endometrial cancer H&E slides in various prediction tasks. The effectiveness of the additional convolutional layer and the integration of patients’ age and BMI through a fourth branch was not found to be significant, however.

### Accurate predictions of histological and molecular subtypes

Panoptes2 achieved a 0.969 (CI: 0.905-1) per-patient level AUROC in classifying samples into endometrioid or serous histological subtypes with an F1 score of 0.75. The precision was 1 and the recall was 0.6 respectively at per-patient level. Panoptes models were in the leading positions followed by InceptionV3 and InceptionV2, all of which had per-patient AUROC above 0.9. For molecular subtyping tasks, we applied all architectures on four binary tasks, each aimed at predicting one molecular subtype versus all others. Panoptes1 achieved a per-patient AUROC of 0.934 (CI: 0.851-1) in predicting CNV-H while all other Panoptes models achieved an AUROC above 0.88, outcompeting the baseline models by 5.8% to 23.3%. The best F1 score was 0.8 with precision of 0.727 and recall of 0.889 respectively. This model also achieved sensitivity of 0.889 and specificity of 0.906 when using 0.5 as the cutoff point of prediction scores. For CNV-L subtype classification, Panoptes1 achieved a per-patient AUROC of 0.889 (CI: 0.755-1), outcompeting the baseline models by 12%. The F1 score was 0.75 with precision of 0.857 and recall of 0.667. For MSI-high, the best per-patient AUROC was 0.827 (CI: 0.705-0.948) and F1 score was 0.615. A POLE subtype classification model achieved per-patient AUROC of 0.681 (CI: 0.499-0.863). However, we improved the POLE subtype classification with a multi-model system introduced in later section.

Although most CNV-H cases are of serous subtype, a portion of high-grade endometrioid cancers are also classified as CNV-H. To further assess whether machine learning models could capture the heterogeneity within this histological subtype, we trained models to predict CNV-H status in endometrioid samples. The Panoptes1 architecture was able to achieve a per-patient AUROC of 0.958 (CI: 0.886-1) and F1 score of 0.667 on this task, suggesting that the model utilized features that were not strongly associated with histological subtype to predict molecular subtype. All Panoptes models also outcompeted baseline models in this task. In addition, we trained models to predict mutation status of 18 driver genes. Panoptes2 was able to predict a *TP53* mutation with a per-patient AUROC of 0.873 (CI: 0.768-0.977) and F1 score of 0.56. *FAT1* mutation was predicted using Panoptes2 (with clinical feature branch) with a per-patient AUROC of 0.835 (CI: 0.666-1) and F1 score of 0.545. Other gene mutations, including *ZFHX3, PTEN, FGFR2, MTOR, CTCF*, and *PIK3R1*, were also predicted with a per-patient AUROC above 0.7. The full statistical metrics for all the prediction models are in Table S1.

### Feature extraction and whole-slide visualization revealed correlations and differences between histological and molecular features

To visualize and evaluate features learned by the models for each task, we extracted the activation maps before the final fully connected layer of the test set tiles. 20000 tiles’ activation maps were then randomly sampled for each task. These activation maps were dimensionally reduced and displayed on 2D tSNE plots, where each dot represents a sampled tile and was colored according to the positive prediction scores (Figure 3). As we expected, tiles were generally clustered by their predicted groups. By replacing dots with the original input tiles of different resolutions, we were able to discover features that correlated with the predictions corresponding to the specific histological or molecular classification task. For example, features of predicted histologically serous and endometrioid were drastically different (Figure 3A). In the cluster with high prediction scores of serous subtypes, we observed typical serous carcinoma features, such as high nuclear grade, papillary growth pattern, elevated mitotic activity, and slit-like spaces. Tiles in the cluster of predicted endometrioid cases showed low nuclear grade, glandular growth pattern, cribriform architecture, and squamous differentiation. Myometrium and other non-tumor tissue tiles were located in the middle of the tSNE plot with prediction scores between 0.4 and 0.6. These observations suggested that our models were able to focus on the tumor regions of H&E slides and make histological subtype predictions based on features that were also recognized by human experts in pathology.

**Figure 3.**
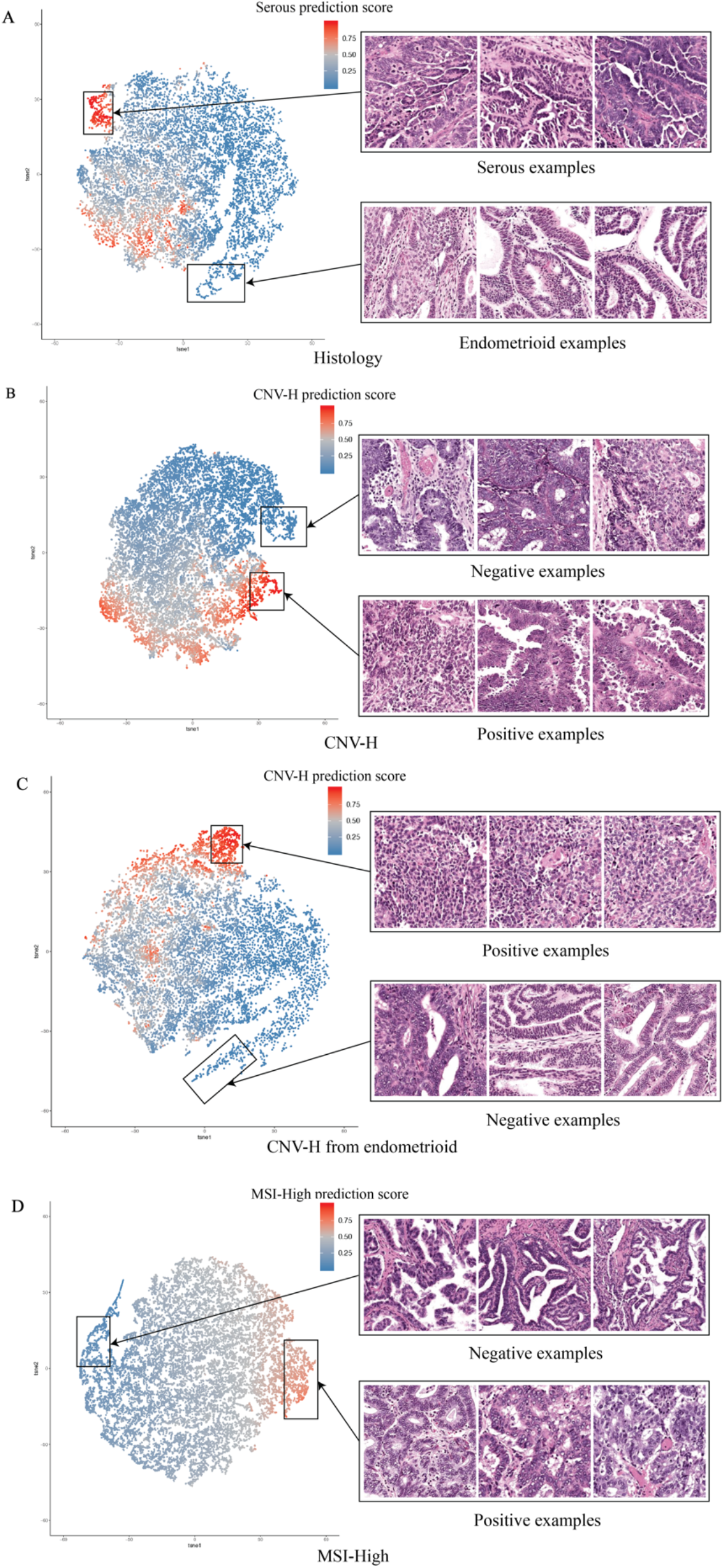
Extraction and visualization of features learned by the models with tSNE. Each point represents a tile and is colored according to its corresponding positive prediction score. (A) Histologically serous and endometrioid features from a Panoptes1 model. (B) CNV-H positive and negative features from a Panoptes4 model. (C) CNV-H positive and negative features in the histologically endometrioid samples from a Panoptes1 model. (D) MSI-High positive and negative features in the histologically endometrioid samples from a Panoptes3 with clinical features model.

The features learned by molecular subtype prediction models were also revealed with the same feature extraction method. We noticed that in the CNV-H prediction model, two distinct subgroups were recognized in the predicted CNV-H cluster, associated with histological serous and high grade endometrioid subtypes, respectively (Figure 3B). The predicted-CNV-H serous tiles mostly showed high nuclear grade, gland formation, and elevated mitotic activity, while the predicted-CNV-H high grade endometrioid tiles exhibited solid growth pattern and focal glandular differentiation. In contrast, in the non-CNV-H cluster, tiles were mostly low-grade endometrioid carcinoma with low nuclear grade, gland formation, and squamous differentiation (Figure 3B). To confirm that the tumor grade was the major factor to distinguish CNV-H molecular subtype in endometrioid samples, we unveiled the features learned by the CNV-H prediction model trained only on endometrioid images (Figure 3C). As we expected, high-grade endometrioid carcinoma tiles were observed mostly in the CNV-H cluster, leaving the low-grade tiles in the non-CNV-H cluster. In both of these CNV-H models, the ambiguous regions were mostly occupied by non-tumor tissue. We also visualized the major pattern learned by the model to distinguish MSI-high subtype images from others (Figure 3D). Tiles in the MSI-H cluster were mostly low grade endometrioid carcinomas with gland formation, tumor infiltrating lymphocytes, and peritumoral lymphocytes, consistent with the observation that heavy mutation load of MSI-high tumors lead to high immunogenicity and a host immune response (Shia et al., 2008; Yamashita et al., 2018).

In addition to the subtypes, patterns related to some mutations were also revealed. A *PTEN*-mutated cluster mostly contained tiles of low grade endometrioid carcinomas with gland formation and low nuclear grade (Figure S4A) while *TP53*-mutated tiles were generally serous carcinomas with high nuclear grade and abundant tufting and budding (Figure S4B). Furthermore, low grade endometrioid carcinoma tiles with gland formation, low nuclear grade, and abundant tumor infiltrating lymphocytes were present in the *ZFHX3*-mutated cluster while those with much less lymphocytes were in the wild type cluster (Figure S4C). High grade endometrioid carcinoma tiles with diffuse solid growth and low nuclear grade were depicted in *FAT1*-mutated cluster while low grade endometrioid carcinomas with gland formation, low nuclear grade, and cribriform architecture were in the wild type cluster (Figure S4D). These findings may result from the correlation between mutation status and histological or molecular subtypes described above, as *PTEN* and *TP53* mutations were mainly found in endometrioid and serous subtypes, respectively, while *ZFHX3* and *FAT1* mutation status showed correlation with the heavily mutated MSI-H and *POLE* molecular subtypes (Figure S1C).

Additionally, we were also interested in visualizing the spatial distribution of features on the whole slide level. Prediction of tiles from the test sets were aggregated back to the size of original slides in the form of heatmaps, where hotter tiles corresponded to higher positive prediction scores. Whole slide visualization revealed that our models tended to have extreme prediction scores on tumor regions instead of non-tumor tissues such as myometrium (Figure S5). The first slide in Figure 4 was from an endometrioid and CNV-H case while the second slide was from a serous and CNV-H case. Models correctly predicted both tasks for the two slides. By comparing the prediction of histological subtypes and CNV-H, we found that the models were focusing on different yet related features in these 2 prediction tasks. In the first slide, the areas predicted to be endometrioid were largely classified as CNV-H while in the second slide, most areas predicted as serous were also classified as CNV-H. This suggested that although most CNV-H samples were histologically serous, our models relied on some additional features other than those of histological subtypes, likely tumor grade, to separate CNV-H samples from endometrioid samples.

**Figure 4.**
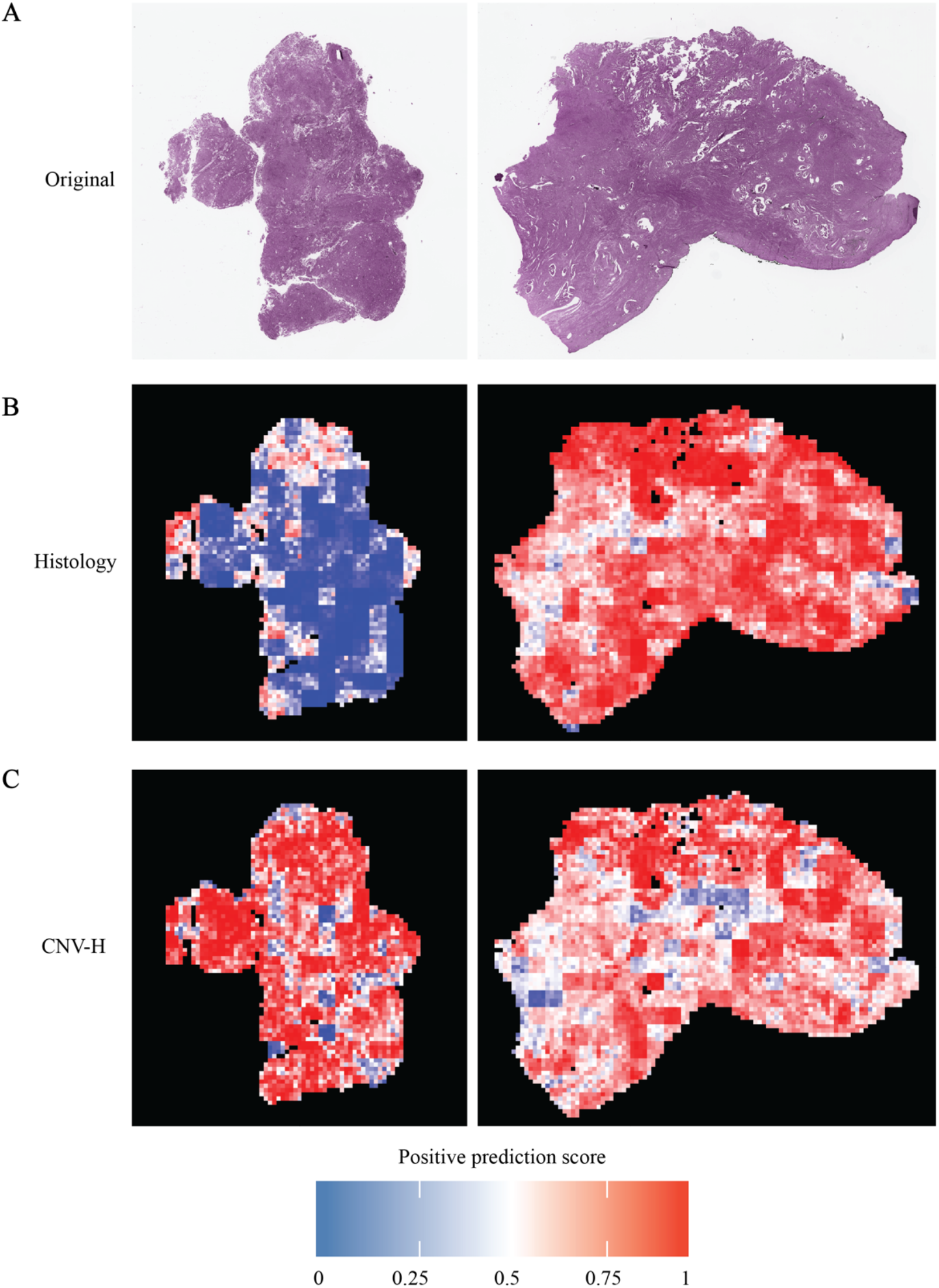
Whole slide predictions showing some features of determining histological subtype and CNV-H are distinct. (A) The first example slide is from a CNV-H but histologically endometrioid case while the second example slide is from a CNV-H and serous tumor. (B) Whole slide histology prediction of examples in (A) from a Panoptes2 model with hotter regions being predicted more of serous while cooler regions were more endometrioid. (C) Whole slide CNV-H prediction of examples in (A) from Panoptes1 (first example) and Panoptes4 (second example) models with hotter regions being predicted more of CNV-H.

### Generalizability and potential clinical capability of the models

To ensure the generalizability of models, especially those with Panoptes architectures, we adopted cohort independent data split in addition to mixed random data split and retrained all predictive models from scratch (Figure 5A, 5B). In the cohort independent data split trials, models were trained and validated only on the data from TCGA at a 9:1 split ratio, and used samples from CPTAC as an independent test set for all the prediction tasks. Therefore, the size of training and validation sets of these trials were smaller and less diversified than the mixed random data split trials. All other hyperparameters remained the same for the training, validation, and testing workflow. The AUROC of CPTAC independent test set indicated that Panoptes-based models still had better performance than the baseline models in general (Figure S2C, S2D). The best performing models based on CPTAC independent test set were compared side-by-side with the best models in mixed random split trials (Figure 5C, 5D). A Panoptes4 model achieved an AUROC of 0.962 (CI: 0.926-0.999) with F1 score of 0.696 at per-patient level in predicting histological subtypes, which were similar to the best model on the mixed random data split. In the CNV-H molecular subtype prediction task, a Panoptes3 model showed an AUROC of 0.87 (CI: 0.753-0.987) with F1 score of 0.667 at per-patient level. Slightly lower performances were also observed in prediction tasks using cohort independent data split at per-patient level, including MSI-high, POLE, *TP53*, and *FAT1*. However, higher statistical metrics were observed in some prediction tasks, such as *PTEN, KRAS, BRCA2*, and *CTNNB1*. Interestingly, even though the per-patient level metrics were lower in the cohort independent data split trials than in the mixed data split trials for some prediction tasks (CNV-H, *TP53, CTCF*), their per-tile level metrics were higher. In addition, we compared Panoptes-based models’ performance side-by-side in mixed random split trials and cohort independent split trials and the results were similar to the best performing models’ comparisons (Figure S6). The full table of statistical metrics of the test set in the cohort independent split trials are in Table S2.

**Figure 5.**
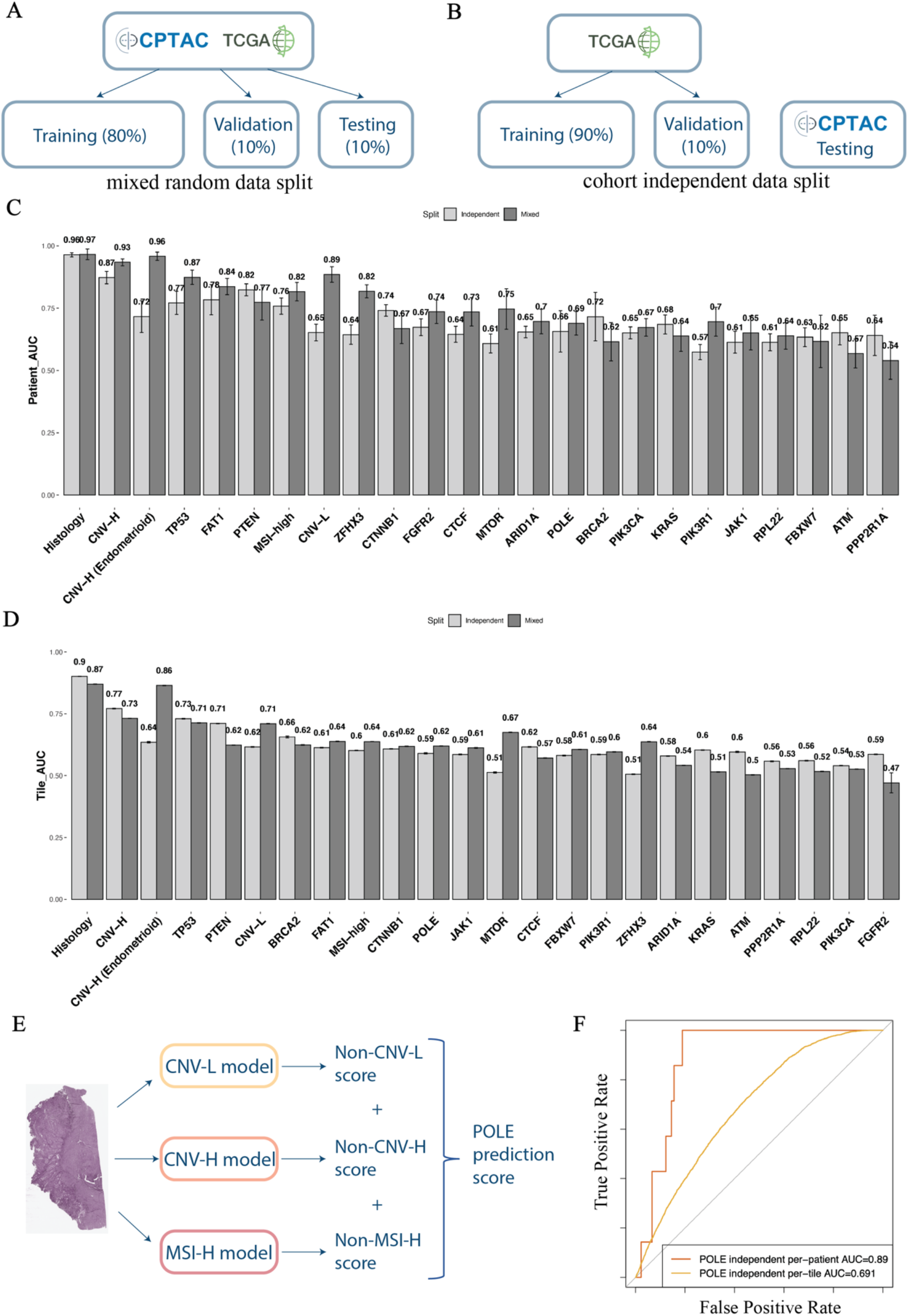
Comparisons of AUROC between the best models in mixed random split trials and cohort independent split trials and multi-model system for better POLE subtype classification. (A) Mixed random data split demonstration. (B) Cohort independent data split demonstration. Per-patient (C) and per-tile (D) level AUROC of the best performing models in each task with mixed random data split (dark) and the cohort independent data split (light). (E) Multi-model system to indirectly predict POLE molecular subtype. (F) ROC curves at per-patient and per-tile level of multi-model POLE classification system.

The best performing model for POLE subtype classification with cohort independent data split achieved per-patient level AUROC of 0.679 (CI: 0.42-0.939), lower than the other three molecular subtypes classification models. To improve the POLE classification, a multi-model system was built by aggregating negative prediction scores of the other three molecular subtypes (Figure 5E). A system consisting of Panoptes2 models of CNV-H and CNV-L and InceptionResnetV1 model of MSI_high with cohort independent data split achieved per-patient AUROC of 0.89 (CI: 0.821-0.96) and per-tile AUROC of 0.691 (CI: 0.683-0.7) for POLE subtype (Figure 5F). The full table of statistical metrics of POLE multi-model classification systems are in Table S3.

Our results demonstrated that the Panoptes model generalize well across cohorts. To further illustrate potential clinical capability, we gathered another independent test set consisting of 137 FFPE H&E slides from 41 patients at NYU hospitals to test the trained models of some prediction tasks, including histological subtypes, CNV-H, CNV-L, MSI-high, and *TP53* mutation (Table S4). This clinical data set was more diversified histologically as it contained not only serous and endometrioid samples, but also samples of rare histological subtypes, including clear cell, carcinosarcoma, mesonephric like, and mixed histology. As genomic sequencing data were not available for this cohort, we used immunohistochemically identified *P53* overexpression as surrogate label for TP53 aberration. The Panoptes2 histological subtype predictive model trained on mixed TCGA-CPTAC dataset achieved a per-patient level AUROC of 0.913 (CI: 0.816-1) on the NYU test set with F1 score of 0.714 (Figure 6A). Notably, this model was the best performing model based on both the mixed held-out test set and the NYU test set. Although the rare histological subtype samples were excluded in statistically metrics calculation, their mean prediction logits generally lay between serous and endometrioid samples, suggesting that they could be linearly separable by setting up appropriate thresholds (Figure 6B). The Panoptes4 CNV-H predictive model, which was the best performing one according to the mixed test set, achieved AUROC of 0.795 (CI: 0.66-0.931) on the NYU test set, lower than 8 other trained models with AUROC ranging from 0.818 to 0.894 (Figure 6C and Table S4). Since there was only 1 endometrioid and CNV-H patient case in the NYU test set, some statistical metrics, such as per-patient level AUROC, could not be calculated for the CNV-H in endometrioid predictive models. The per-tile level AUROC of the Panoptes2 model on NYU test set was 0.919 (CI: 0.911-0.926), similar to its performance on the mixed test set (Table S1 and Table S4). The Panoptes1 CNV-L predictive model achieved per-patient AUROC of 0.85 (CI: 0.732-0.968) on the NYU test set, similar to its performance on the mixed test set, which was the best among all models (Figure 6D and Table S1). The best performing MSI-high predictive model on mixed test set was also the best one on NYU test set (Figure 6E). Interestingly, the Panoptes2 *TP53* aberration prediction model achieved a higher per-patient level AUROC on NYU test set (0.92; CI: 0.836-1) than on mixed test set (Figure 6F). This was likely due to the fact that *TP53* aberration determined at expression level were easier for the model to detect, further supporting our conclusion that the model recognized morphological manifestations of genetic aberration. Generally, the performance of TCGA-CPTAC mixed data split trained models on NYU test set were comparable to their performance on mixed held-out test set in these tasks, suggesting that these models are generalizable on independent clinical samples. We also tested models training only on TCGA samples (cohort independent data split) on NYU test set, and the performance are mostly slightly lower than the mixed data split trained ones (Table S4).

**Figure 6.**
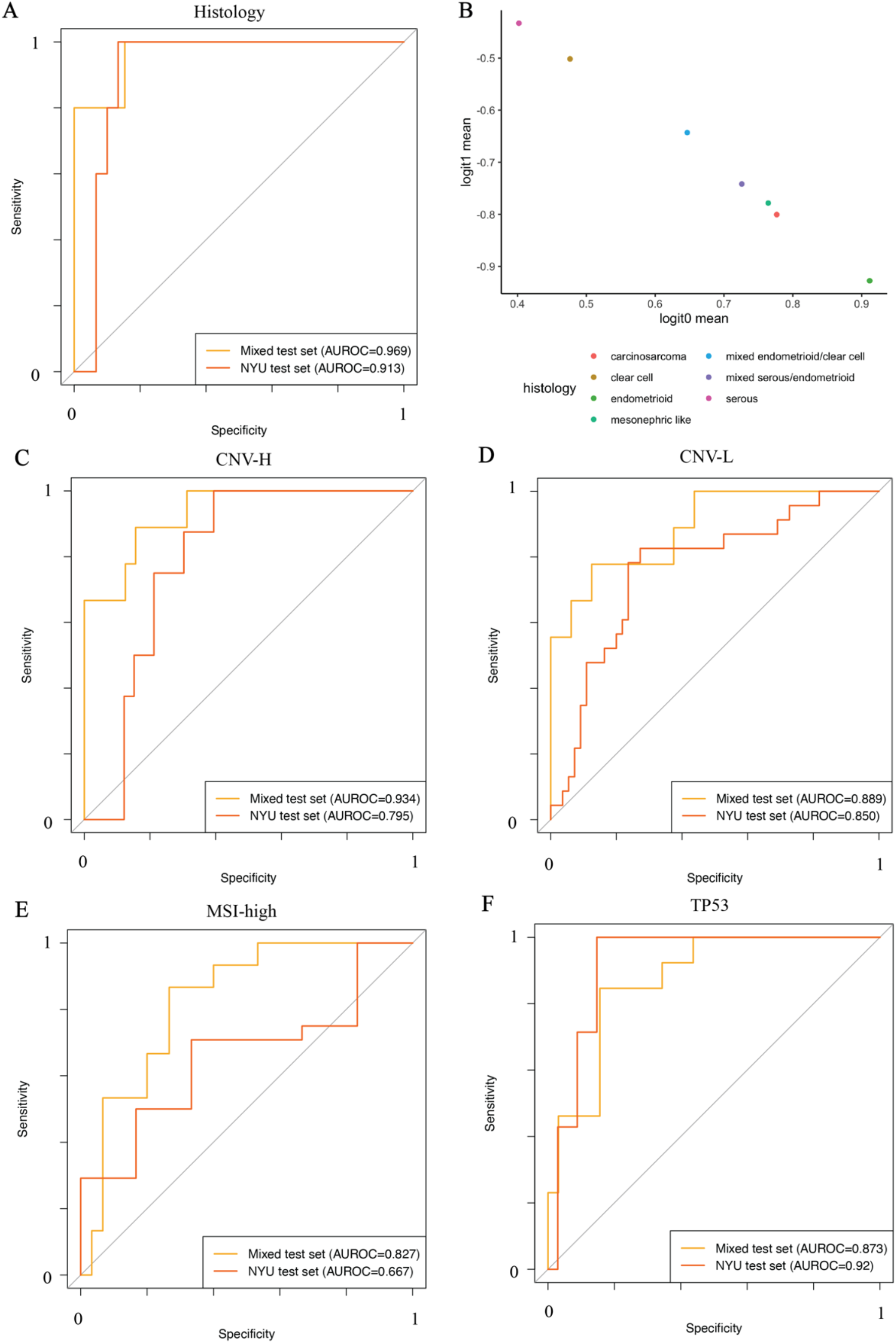
ROC curves and AUROC of the best trained models of key tasks showed promising predictive power on the independent clinical dataset. (A) Trained Panoptes2 histological subtype predictive model on the mixed TCGA and CPTAC held-out test set and the NYU test set. (B) Mean prediction logits by histology from histological subtype predictive model of NYU test set samples. (C) Trained Panoptes4 CNV-H subtype predictive model on the mixed TCGA and CPTAC held-out test set and the NYU test set. (D) Trained Panoptes1 CNV-L subtype predictive model on the mixed TCGA and CPTAC held-out test set and the NYU test set. (E) Trained InceptionResnetV1 MSI-high subtype predictive model on the mixed TCGA and CPTAC held-out test set and the NYU test set. (F) Trained Panoptes2 *TP53* mutation predictive model on the mixed TCGA and CPTAC held-out test set and the NYU test set.

## Discussion

Our study introduced a novel multi-resolution InceptionResnet-based convolutional neural network architecture, Panoptes, which was able to predict endometrial cancer histological and molecular subtypes as well as mutation status of critical genes based on H&E slides and generalized well on independent test sets. The AUROC of classifying endometrioid and serous histological subtypes by our best architecture model was 0.969 (CI: 0.905-1). Moreover, the models can distinguish the most lethal molecular subtype, CNV-H, with exceptionally high accuracy (AUROC 0.934). It is worth noting that our models can also precisely identify the CNV-H samples from a histologically endometrioid carcinoma (AUROC 0.958), which is one of the more controversial and complex patient subgroups in endometrial cancer subtyping. In addition to the CNV-H, we were also able to predict other molecular features with acceptable performance, which are currently not possible for pathologists to determine without ancillary studies, such as sequencing or immunohistochemistry. These include CNV-L molecular subtype (AUROC 0.889), the DNA-mismatch repair deficiency-related MSI-high molecular subtype (AUROC 0.827), the mutation of the CNV-H signature gene *TP53* (AUROC 0.873) as well as *PTEN* (AUROC 0.781), *FAT1* (AUROC 0.835), and *ZFHX3* (AUROC 0.824). Although the direct predictive model for POLE molecular subtype did not achieve promising results due to insufficient training samples, an indirect multi-model system approach allowed us to classify POLE subtype effectively (AUROC 0.89). Statistical analyses proved the success of our prediction tasks. In addition, we tested and showed that our multi-resolution Panoptes-based models performed significantly better than InceptionResnet-based models in most of our prediction tasks. We implemented two modifications to Panoptes, including an additional convolutional layer and integration of clinical features, but were not able to observe significant improvement in performance for majority of the tasks. By extracting and clustering abstract representation of tiles constructed by the model, we discovered critical features to distinguish subtypes and mutations, particularly the tumor grade in determining CNV-H cases from non-CNV-H samples. We justified the generalizability and potential clinical applicability of our models by testing on an independent dataset of samples from NYU hospitals for some most promising prediction tasks, including histological subtypes, CNV-H, CNV-L, MSI-high, and *TP53* mutation. Even though the NYU test set contained some samples of rare histological subtypes, their prediction logits suggested that they could potentially be separated by our models with additional simple thresholds. Moreover, we observed a noticeable drop of performance for MSI-high prediction model on NYU test set, which is similar to the level of drop of performance in previous studies when applying models to independent clinical images (Fu et al., 2020; Kather et al., 2020). We believe that more MSI-high training samples would be essential to allow the model to be more generalizable to clinical images. We were not able to conduct generalizability tests for all prediction tasks using this NYU deidentified clinical dataset due to lack of relevant sequencing information. Instead, for other prediction task, including the multi-model system for POLE, we retrained the models with only TCGA samples and tested on CPTAC samples to prove their generalizability. Although slightly lower performances were observed in the CPTAC-only testing trials for some prediction tasks, we believe that it was most likely caused by smaller and less diversified TCGA-only training set. The differences in feature distributions among samples in the training set and the independent test sets could also be factors that affected the models’ performance.

Examining H&E slides is still currently the most widely used techniques for pathologists to confirm endometrial cancer histological subtypes in the clinical setting. Our models showed great potential in assisting pathologists making decisions and improving diagnostic accuracy. Given most H&E slides can be tiled into less than 5000 tile-sets (Figure S1E, S1F, S1G), with a processing speed of 22 tile-sets per second (1310 tile-sets per minute) on a Quadro P6000 GPU, our models can analyze a slide within 4 minutes. This means that these models can work simultaneously with pathologists to serve as references. We have shown that the model utilized human interpretable features to perform histological and molecular classification tasks. With whole slide visualization, the reassembled per tile predictions can provide a thorough examination of the H&E slide and a detailed layer containing potential hotspot features, which may also include regions that could possibly be neglected by pathologists. However, due to the time-consuming H&E slide scanning and tiling processes, multiple optimizations need to be implemented before the system could be deployed in practice.

Both histological and molecular features’ labels of TCGA and CPTAC samples have been validated by many scientists and clinicians before and after the publication of their studies. The NYU test set labels were also validated by pathologists. However, as tile labels were assigned at per-patient level, within-slide heterogeneity would still lead to noise in the true labels, such that features in a local region may not match the characteristics of the assigned classification. Therefore, we believe that the per-patient metrics are more accurate than per-tile metrics in terms of assessing a model’s performance. The performance can be further improved if more detailed annotations existed on the slides. From the visualization results, we noticed that our models were more likely to give non-tumor tissue tiles ambiguous prediction scores (0.4-0.6). Therefore, building a segmentation model or set up a threshold to exclude these irrelevant non-tumor tissue, such as myometrium, may also significantly enhance the overall performance of our models. Although the TCGA, CPTAC, and NYU datasets cover a variety of endometrial carcinoma samples, it may not reflect the full pathological diversity and feature distribution of endometrial cancer. More diversified training sets could improve the robustness of the models and ideally boost the performance in prediction.

Overall, we demonstrated that our multi-resolution convolutional neural network architecture, Panoptes, can be a practical tool to assist pathologists classifying endometrial cancer histological subtypes and, more importantly, to provide additional information about patients’ molecular subtypes and mutation status in a much more rapid fashion and without the need for sequencing. In addition to per-patient level prediction, the model would also be able to highlight regions with human interpretable features on the slide. Moreover, it remains possible that our models have learned visual patterns correlating with molecular features that were not previously annotated by human experts and, thus, requires further investigation. From another perspective, these novel patterns from the H&E slides may be incorporated into the current standards of histological pathology and contribute to improved prognosis and treatment of endometrial carcinoma in the future.

Our future plan includes refining the Panoptes architecture, particularly to determine an effective way to integrate clinical features into the imaging prediction branches to improve the overall performance. Quantification of features could also be added to the Panoptes. Although our statistical analyses demonstrated promising results for various tasks, the models need to be trained on a more diversified data to meet the more stringent criteria for real world clinical application. Therefore, we would work on training our existing models with slides labeled with more details and new datasets that cover more heterogeneity of endometrial cancer in order to make the models more robust and generalizable. Predicting other molecular subtypes and mutations, such as *ARID1A, CTNNB1*, and *JAK1*, which did not have a well-performing model in this study, will be possible once more data are available. In addition to the currently available user interface of Panoptes, we plan to develop a more advanced Graphical User Interface (GUI) that includes all the trained models and outputs visualization and prediction in a fast and user-friendly way, which we are hoping to be deployed and tested in a pathologist’s clinical practice. We would try to train Panoptes-based models to predict features in other types of cancers, such as glioblastoma, melanoma, and lung carcinoma, and it would be very interesting to see how Panoptes performs and what features it captures in these new tasks.

## Acknowledgements

We would like to thank the BigPurple high performance computing team at NYU Langone Health for providing guidance, computational resources, and troubleshooting for this project. We would like to thank the Experimental Pathology Research Laboratory at NYU Langone Health for scanning the H&E slides from NYU hospitals for this project. We would like also thank the high-performance computing team at New York University who maintains the high-performance computing resources at NYU main campus. Special thanks to everyone in the Clinical Proteomic Tumor Analysis Consortium (CPTAC) for generating and maintaining the data we used and previous studies they have done.

## Author Contributions

R.H. designed, trained, and tested the models and wrote most of the codes for this project. W.L. made the feature annotation figure and wrote some of the codes for feature visualization. D.D. reviewed the clinical and histopathological features and coordinated the acquisition of samples from NYU hospitals. N.R. provided model evaluation and improvement ideas. D.F. offered guidance and coordinated the data retrieval and resource allocation. R.H., W.L., and D.D. wrote the paper and N.R. and D.F. edited.

## STAR Methods

### RESOURCE AVAILABILITY

#### Lead Contact

Further information and requests for resources should be directed to and will be fulfilled by the Lead Contact, Runyu Hong.

#### Materials Availability

This study did not generate new unique reagents.

#### Data and Code Availability

The analytic codes are available on GitHub in the following link: https://github.com/rhong3/CPTAC-UCEC. The Panoptes codes with user interface are available on Github in the following link: https://github.com/rhong3/Panoptes. Trained models may be shared upon request. 392 diagnostic slides from 361 Uterine Corpus Endometrial Carcinoma (UCEC) patients in TCGA cohort were downloaded from the NCI-GDC Data Portal. These samples were published in the TCGA pan-cancer atlas. Demographic, genomic, and other clinical features associated with these samples were downloaded from the cBioPortal and the original TCGA UCEC paper supplements (Getz et al., 2013). 107 diagnostic slides from 98 Uterine Corpus Endometrial Carcinoma (UCEC) patients in CPTAC cohort were downloaded from The Cancer Imaging Archive (TCIA). Demographic, genomic, and other clinical features of these patients were published in the CPTAC UCEC paper (Dou et al., 2020). The composition of patients with different features of interests are shown in Figure 1A. Most of the patients in our cohort have only 1 diagnostic slide (Figure S1B). Deidentified digital H&E samples from NYU hospitals used in this study may be released upon request with proper documentations.

## METHOD DETAILS

### H&E Images Preparation

Digital histopathologic images were in SVS or SCN format, which were tuples of the same images with multiple different resolution levels. Slides from the TCGA cohort were scanned with a maximum resolution of 40x while those from the CPTAC cohort and NYU were at 20x maximum resolution. A Python package, Openslide, was used to maneuver the SVS and SCN files. Due to the extremely large size of these images (Figure S1D), they were cut into small tiles in order to be fed into the training pipeline. Multi-threading was used to accelerate this process. Tiles were cut at 10x, 5x, and 2.5x equivalent resolutions and algorithm was used to exclude tiles with more than 40% pixels of white background and irrelevant contaminants (Figure S1E, S1F, S1G). Stain colors of the useful tiles were normalized using the Vahadane’s method during this process (Vahadane et al., 2016). For each of the tasks, the labels were one-hot encoded at per-tile level. The datasets were separated into training, validation, and testing sets at per-patient level with a ratio of 8:1:1 for mixed data split trials. To take advantage of the Tensorflow API and accelerate the training and testing process, tiles were loaded and saved into a single TFrecords file for each set.

### Baseline Models

InceptionV1, InceptionV2, InceptionV3, InceptionResnetV1, and InceptionResnetV2 architecture were trained from scratch and used as the baseline models. InceptionResnets are enhanced architectures of Inceptions with residual connections and a previous study has shown that they are performed generally better than Inceptions in imaging prediction tasks (Szegedy et al., 2017). The auxiliary classifiers of these architectures were opened. We did not modify any part of the backbone of these architectures. Tiles with 10x resolution were input and we used back-propagation, softmax cross entropy loss weighed by training data composition, and Adam optimization algorithm in the training workflow. Here, each single tile image with a label was considered 1 sample. Batch sizes were set to 64 with an initial learning rate of 0.0001 and a drop-out keep rate of 0.3. We tested multiple combinations of hyperparameters and found that this one achieved optimal results for most tasks. The training jobs were run with no fixed epoch number. 100 batches of validation were carried out every 1000 iterations of training and when the training loss achieved a new minimum value after 30000 iterations of training. If the mean of these 100-batch validation loss achieved minimum, the model was saved as the temporary best performing model. The training process stopped when the validation loss did not decrease for at least 10000 iterations. This stopping criterion was only initiated after 100000 iterations of training.

### Panoptes Models

We used 4 different Panoptes architectures with and without the integration of patients’ BMI and age in a fourth branch. Panoptes1 has 3 branches based on InceptionResnet1 and Panoptes2 has 3 branches based on InceptionResnet2. The major difference between Panoptes3 and Panoptes1 and between Panoptes4 and Panoptes2 is the additional 1-by-1 convolutional layer between the concatenation of branches and the global average pooling. All of our Panoptes architectures were trained with randomly initialized network parameters with auxiliary classifiers opened on each branch. Unlike the baseline models, tiles of 10x, 5x, and 2.5x resolutions of the same region on the H&E slide with label were paired and considered as 1 sample as only 1 prediction score was associated with a multi-resolution matrix. Batch size was set to 24, which was the largest number that could fit in the memory of our GPUs. Optimization algorithm, weighted loss function, and other hyperparameters were the same as the baselines. In addition, we applied the same validation method to pick the best performing models and kept the same stopping criterion as the baselines.

### Feature Visualization Based on Tiles

For models with per-patient level AUROC above 0.75 of the test set, we randomly sampled 20000 tiles (tile sets for Panoptes) together with their feature maps before the last fully connected layer in the model, in which each tile or tile set is represented as a 1-dimensional vector. We then used tSNE with initial dimensions of 100 to reduce these 20000 vectors into 2-dimensional space where each point represents a tile or tile set. Generally, points clustered according to their predicted class. By replacing the points on tSNE plots with the original tiles, the features learned by the model for each of the specific class can be observed. We asked experienced pathologists to summarize the typical histological features in each of these clusters.

### Whole Slide Prediction

We built an implementation pipeline that could apply trained models to whole H&E slides and output predictions as heatmaps. The heatmaps could be overlaid on the original slides, which showed the prediction results of different areas. The maximum prediction resolution (each cell of the heatmap) is 299 by 299 pixel at 10x resolution level. Depending on the size of the H&E slides, the time of predicting an intact H&E slides can range from 2 to 40 minutes. The average speed of prediction with Panoptes models is 22 tile-sets per second, or 1310 tile-sets per minute.

## QUANTIFICATION AND STATISTICAL ANALYSIS

The performance was evaluated by applying the trained models to the test set. Each of the classification tasks has its own test set, which consists of slides from patients that had not been in the training or validation sets. Evaluation was performed at both per-patient level and per-tile level. Per-patient level metrics were obtained by taking the mean of all tiles’ metrics that belonged to the same patient. For Panoptes model, a 3-multi-resolution-tile matrix is considered as 1 tile for statistical analyses. Receiver Operating Characteristic (ROC) curve, plotting true positive rate against false positive rate, and the area under the ROC curve (AUROC) were the major factors in evaluation. In addition, Precision Recall Curve (PRC), as well as average precision score (AUPR score), were used to determine the trade-off between false negative rate and false positive rate. We also used accuracy with softmax prediction score directly from the models. If the prediction score was greater than 0.5, it was counted as a positively predicted case. 95% Confidence intervals (CI) of AUROC, AUPR, and accuracy were estimated by the bootstrap method. Other statistical metrics, including sensitivity, specificity, precision, recall, F1 score, etc., were also generated and referred to evaluate the predictive models’ performance. To further validate the effectiveness of the classification models, we did 1-tail Wilcoxon tests between positive and negative tiles in the test sets for each of the tasks. In order to compare performance between Panoptes models and the baselines, for each of the tasks with a patient level AUROC score greater than 0.75, we bootstrapped 50 times at an 80% sampling rate at both patient and tile level and calculated the AUROC for each of these sampled sets. Then, an unpaired 1-tail t-test between the AUROC of Panoptes and its corresponding baseline model was performed. We performed a similar t-test between Panoptes with and without the additional convolutional layer as well as between Panoptes with and without the fourth branch of patients’ BMI and age. Statistical analyses and plotting codes were written in R3.6 and Python3.

## ADDITIONAL RESOURCES

The Panoptes Python3 package version is on PyPI with the following link: https://pypi.org/project/panoptes-he/.

## KEY RESOURCES TABLE

| REAGENT or RESOURCE | SOURCE | IDENTIFIER |
| --- | --- | --- |
| Software and Algorithms |  |  |
| Tensorflow | Abadi et al., 2015 | https://www.tensorflow.org |
| Inception | Szegedy et al., 2015 | N/A |
| InceptionResNet | Szegedy et al., 2017 | N/A |
| Keras | Chollet et al., 2015 | https://keras.io |
| Panoptes | This paper | https://github.com/rhong3/Panoptes |
| cBioPortal | Cerami et al., 2012; Gao et al., 2013 | https://www.cbioportal.org |
| Other |  |  |
| NVIDIA Tesla V100 GPU | NYU Langone Health BigPurple HPC Cluster | http://bigpurple-ws.nyumc.org/wiki/index.php/BigPurple_HPC_Cluster |
| NVIDIA Tesla P40 GPU | NYU HPC Cluster | https://wikis.nyu.edu/display/NYUHPC/High+Performance+Computing+at+NYU |

## Supplementary figures

**Figure S1.**
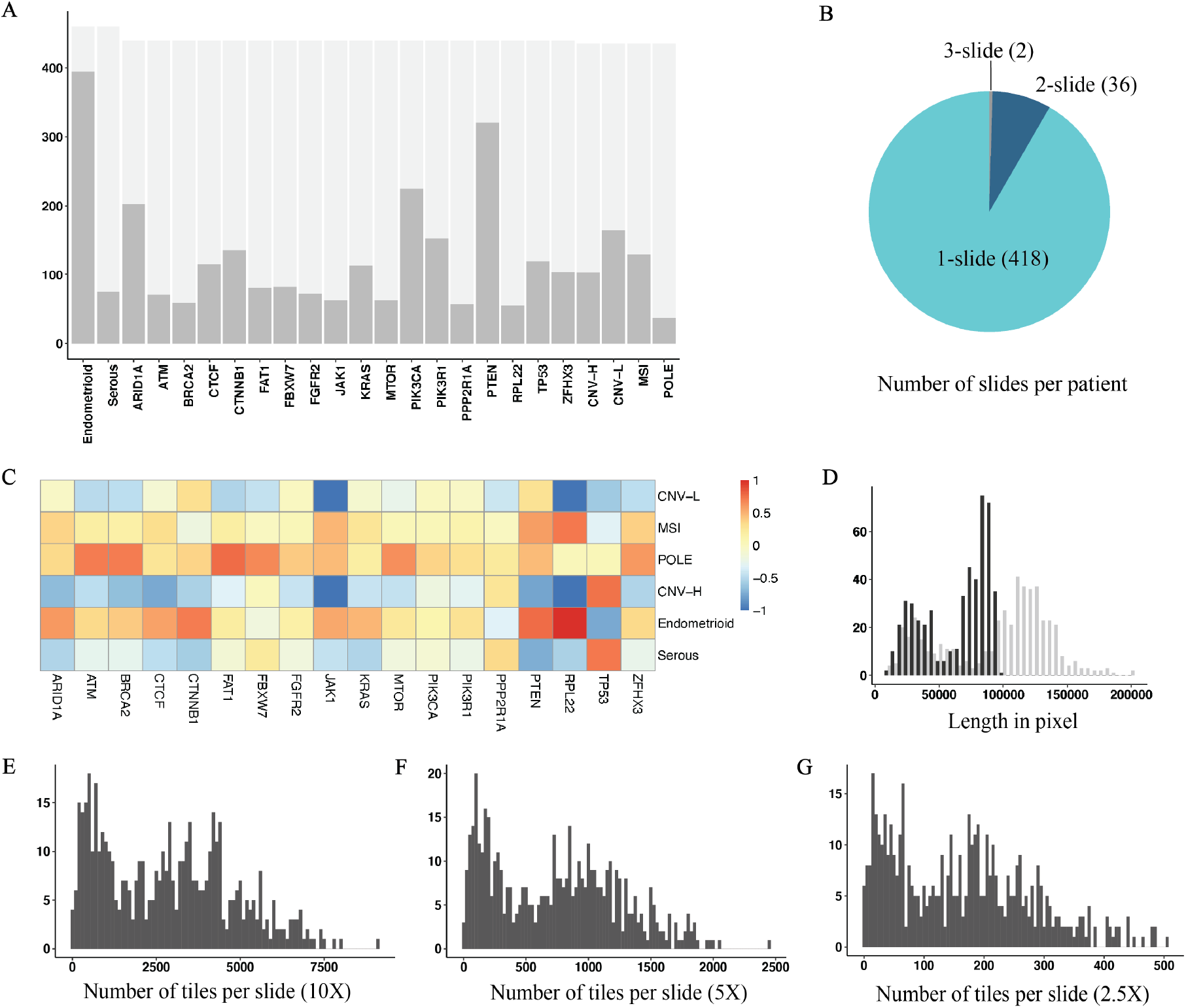
Data summary. (A) Number of patients and composition of true labels in each task. (B) Number of slides per patient in the cohort. (C) Coefficient of colligation between subtypes and mutations. (D) Dimensions of slides in pixel (black: height; grey: width). (E, F, G) Number of tiles per slide at 10X (E), 5X (F), and 2.5X (G) equivalent resolution.

**Figure S2.**
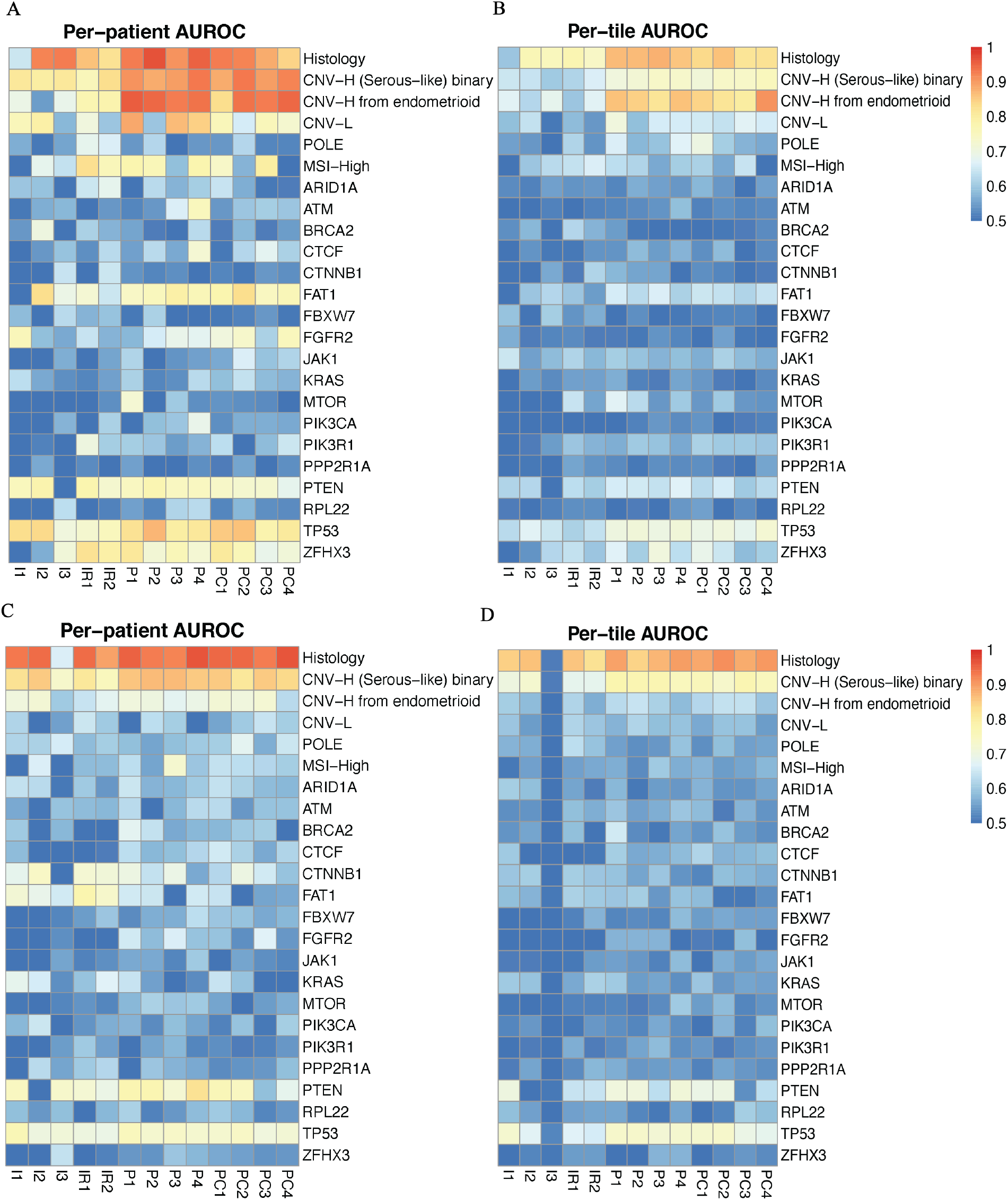
AUROC achieved for Panoptes and baseline models on each prediction task using mixed random data split (A, B) and cohort independent data split (C, D) at per-patient and per-tile level. P represents Panoptes, PC represents Panoptes with clinical features, I represents Inception, and IR represents InceptionResnet.

**Figure S3.**
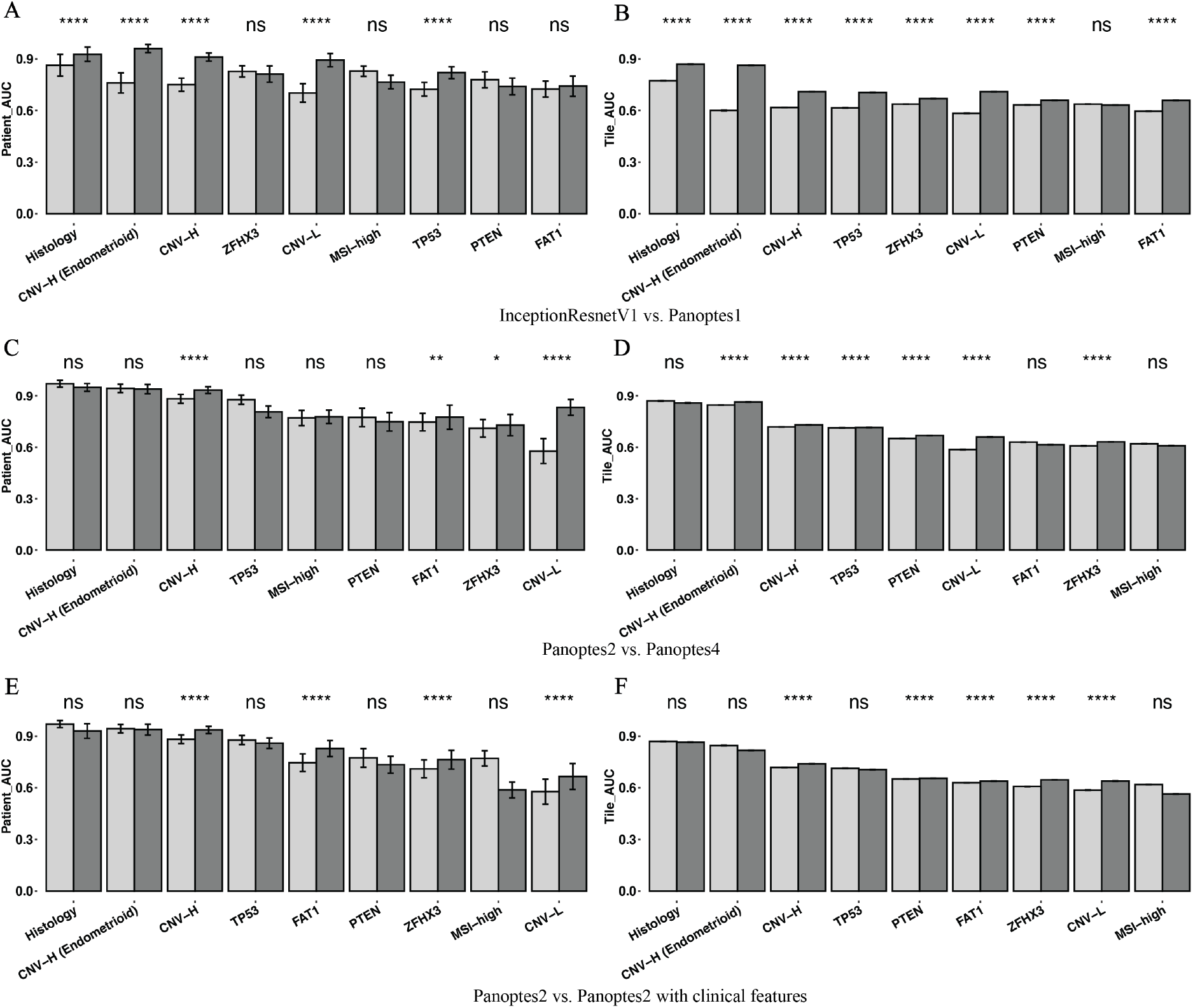
Comparisons of AUROC between architectures on the top eight prediction tasks. (A, B) 1-tail t-test of per-patient (A) and per-tile (B) AUROC between InceptionResnetV1 (light) and Panoptes1 (dark) of the top nine tasks. (C, D) 1-tail t-test of per-patient (C) and per-tile (D) AUROC of Panoptes2 (light) and Panoptes4 (dark) of top nine tasks. (E, F) Bootstrapped per-patient (E) and per-tile (F) 1-tail t-test of AUROC of Panoptes2 (light) and Panoptes2 with clinical features (dark) of top nine tasks.

**Figure S4.**
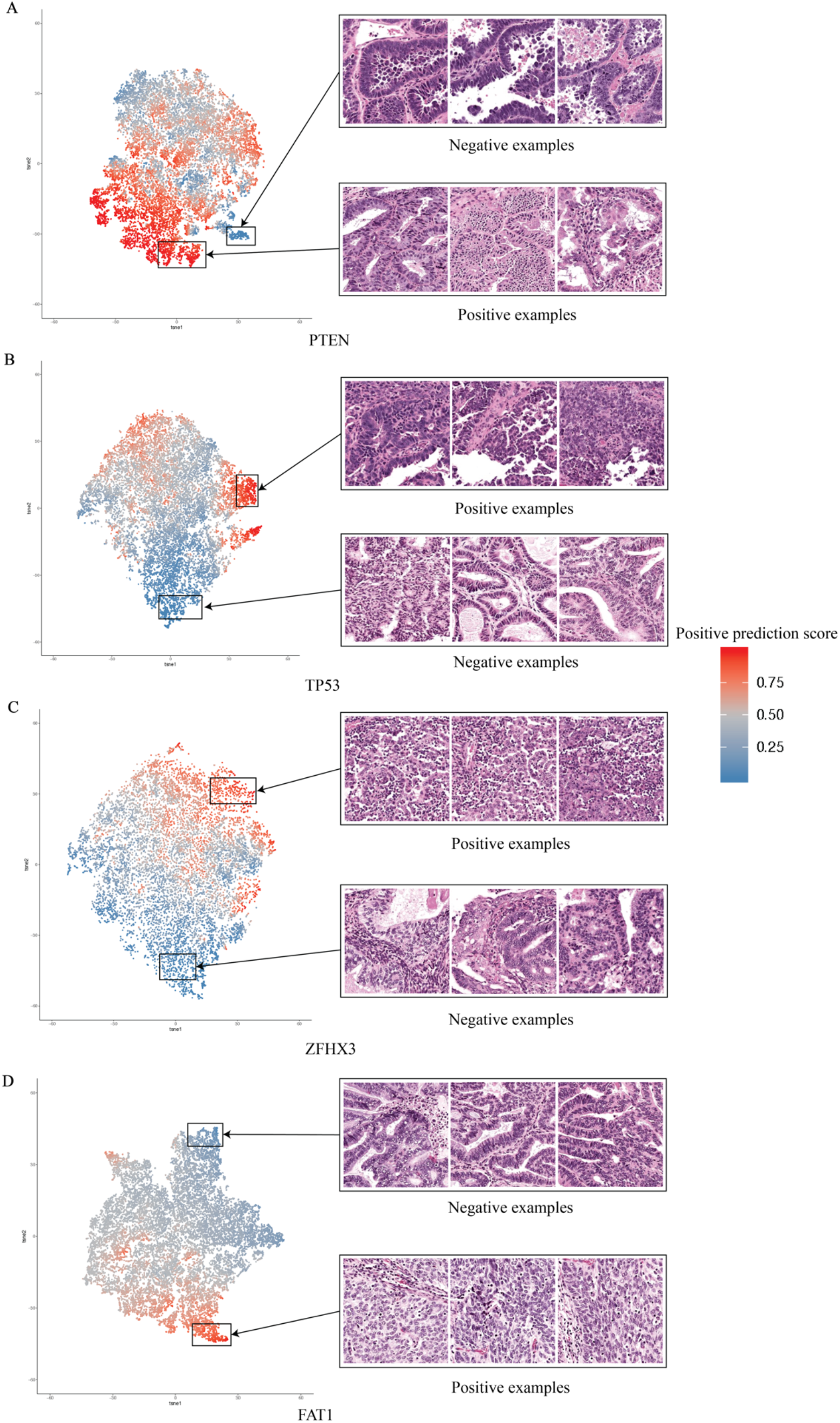
Extraction and visualization of features learned by the models with tSNE. Each point represents a tile and is colored according to its corresponding positive prediction score. (A) *PTEN* from a Panoptes2 model. (B) *TP53* from a Panoptes2 model. (C) *ZFHX3* from a Panoptes1 model. (D) *FAT1* from a Panoptes2 with clinical features model.

**Figure S5.**
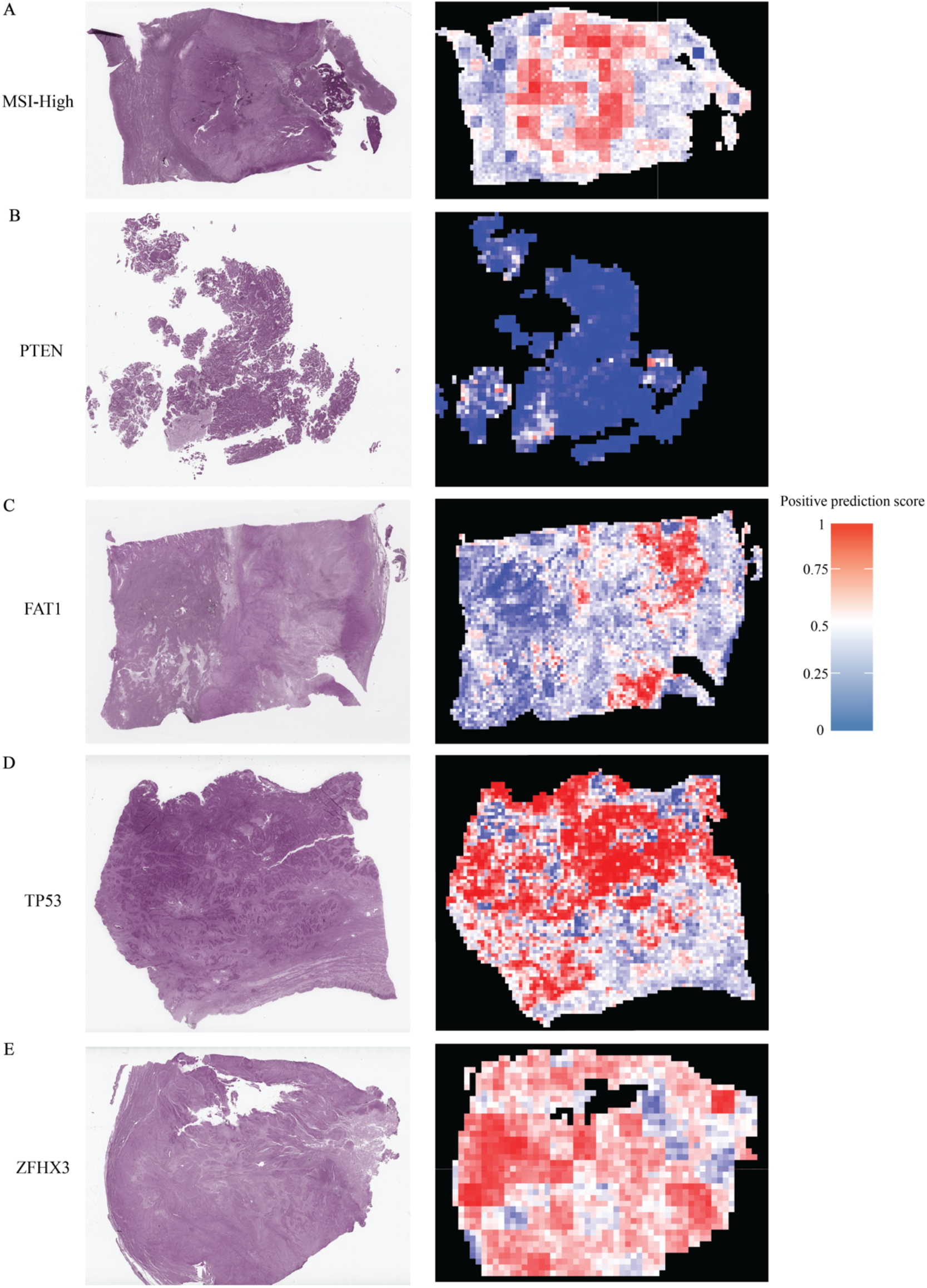
Whole slide predictions with color representing positive prediction scores. (A) Slide from an MSI-High (positive) patient using a Panoptes1 model. (B) Slide from a *PTEN* wild-type (negative) patient using a Panoptes2 model. (C) Slide from a *FAT1* mutated (positive) patient using a Panoptes3 model. (D) Slide from a *TP53* mutated (positive) patient using a Panoptes2 model. (E) Slide from a *ZFHX3* mutated (positive) patient using a Panoptes1 model.

**Figure S6.**
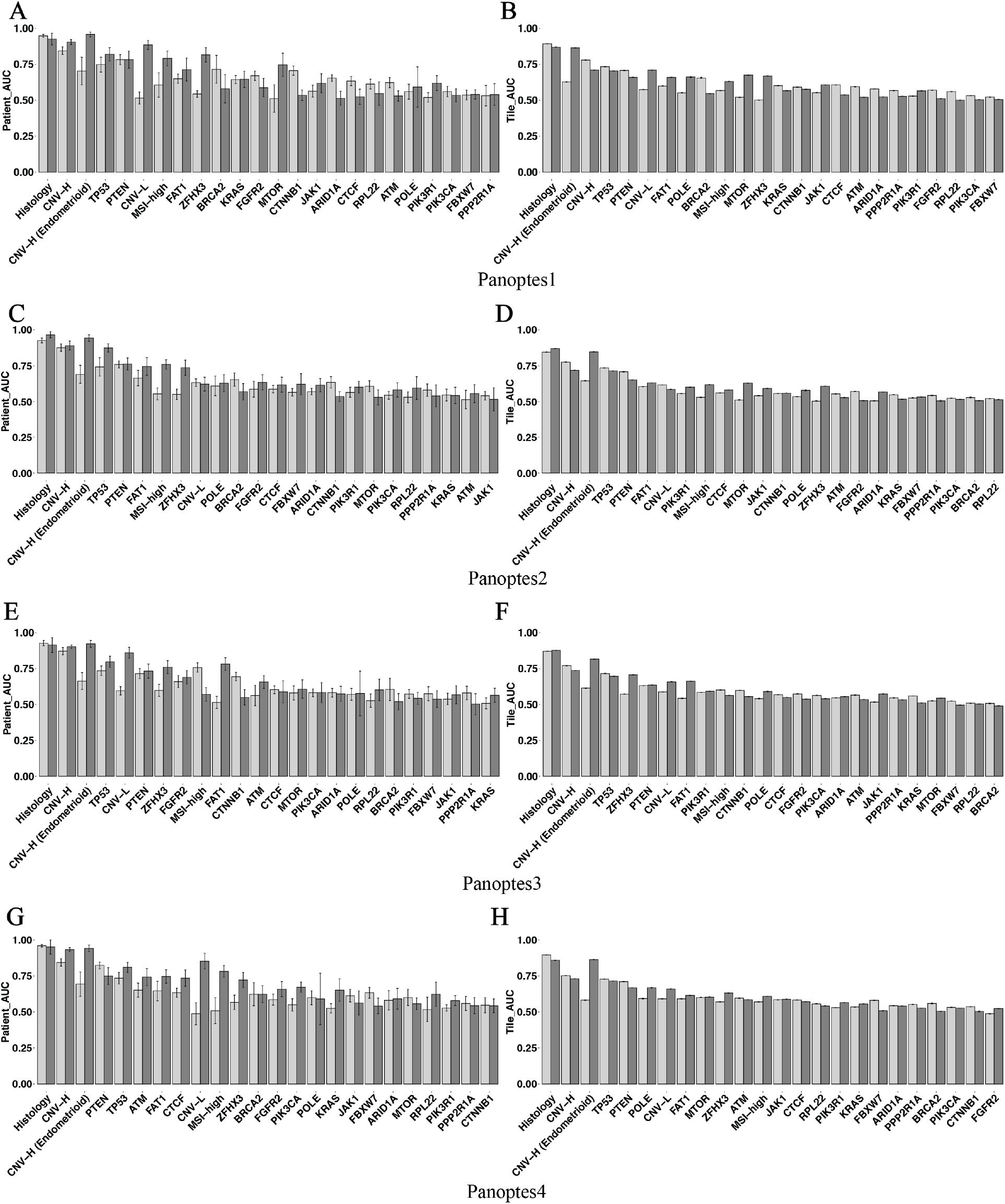
Comparisons of AUROC between the Panoptes models in mixed random split trials and cohort independent split trials. Per-patient and per-tile level AUROC of Panoptes1 (A, B), Panoptes2 (C, D), Panoptes3 (E, F), and Panoptes4 (G, H) models in each task with mixed random data split (dark) and the cohort independent data split (light).

